# Glutamatergic system in the pelagic tunicates

**DOI:** 10.64898/2026.09.02.748133

**Authors:** Leonid L. Moroz, Tigran P. Norekian

**Author notes:** Corresponding author Leonid L. Moroz. both authors contributed equally.

## Abstract

Tunicates are a sister lineage to vertebrates, with compact, relatively simple nervous systems featuring a single central ganglion, reflecting a minimal complement of chordate functional architecture. Although glutamatergic neurons are the most abundant population in vertebrates, their ancestry remains unclear. Here, we used glutamate immunohistochemical labeling (Glutamate IR) to identify glutamatergic elements in the neural system of the pelagic tunicate *Doliolum* sp. (Thaliacea). Glutamate IR was observed in all major nerves of the central ganglion, including motor-like terminals on the circular bundles of swim muscles, which were themselves labeled. However, the neuronal somata in the central ganglion were not labeled, suggesting glutamate accumulation in axonal processes and terminals. In contrast, we did not identify GABA-containing neural elements. This study suggests that glutamatergic systems were elaborated in the common ancestor of tunicates and vertebrates, although the functional role of glutamate and its role in muscular control need further investigation in these pelagic tunicates.

## 1. INTRODUCTION

The origin and early diversification of neural systems in chordates, and their relationships to other animal phyla, including two basal deuterostome groups with predominantly diffuse nerve nets (Hemichordata and Echinodermata), remain elusive. Three major evolutionary trajectories (subphyla) within Chordata are represented by cephalochordates, tunicates, and vertebrates; each has a centralized nervous system, either as a dorsal tube (e.g., amphioxus and vertebrates) or a central ganglion (tunicates). The central ganglion may represent a secondary simplification of the adult neural system after metamorphosis, as extensively studied in ascidians, which lose the larval dorsal cord (Manni, Pennati, 2016).

Pelagic tunicates (class Thaliacea), with middle Cambrian ancestry (Zeng et al. 2026), comprising pyrosomes (order Pyrosomatida), doliolids (Doliolida), and salps (Salpida), have received less attention from neuroscientists because they inhabit the open ocean and are less accessible for experimental studies, making them difficult to maintain in the laboratory. They remain a biological enigma in numerous aspects of animal organization (Piette, Lemaire, 2015).

Thaliaceans are metabolically active, oxygen-demanding and move through the water by contracting and pumping water through their gelatinous bodies (Bone and Trueman, 1983; Madin, 1990; Damian-Serrano et al., 2025), feeding on plankton as individuals or in colonies (Bone, 1998). They are known for their complex alternation of sexual and asexual generations. Sexually produced individuals reproduce asexually by budding a group of blastozooids (i.e., zooids derived from buds), which are usually released as aggregates. Each member of the aggregate, in turn, reproduces sexually, typically producing a single egg that is fertilized internally and develops into a small adult oozooid (i.e., derived from an egg) within the parent (Bone, 1998).

Classical studies by several prominent investigators provided an overview of Thaliaceans’ neural systems (Fedele, 1923, 1938; Bone, 1959; Bulloch and Horridge, 1965; Manni, Pennati, 2016). Their CNS consists of a small dorsal ganglion, the only concentrated neural center. The ganglion is spherical, with a central neuropile and a couple of surrounding layers of about one thousand neuronal cell bodies with an eye-like structure at the dorsal side (Fedele, 1923, 1938). A series of radiating peripheral nerves extends from the dorsal ganglion to all other parts of the barrel-shaped body. However, knowledge of the neurochemical architecture of any Thaliacea is limited. For example, serotonin immunolabeling revealed a broad topographical diversity of neuronal and non-neuronal (endostyle) cell types of unknown function (Pennati et al., 2012; Valero-Gracia et al., 2016). No other information is available on neurotransmitter chemistry in pelagic tunicates.

In this study, we focus on the distribution of putative glutamatergic neurons in an open-water Pacific doliolid species (**Fig. 1**) with a habitat range from Alaska to South America. Glutamate is the most abundant neurotransmitter in vertebrate brains, representing more than 60% of the total neuronal population, but only about 10% in cephalochordates (Candiani et al., 2012) and in benthic tunicates, such as the widely used reference genus *Ciona* (Takamura et al. 2010; Manni, and Pennati, 2016).

**Figure 1.**
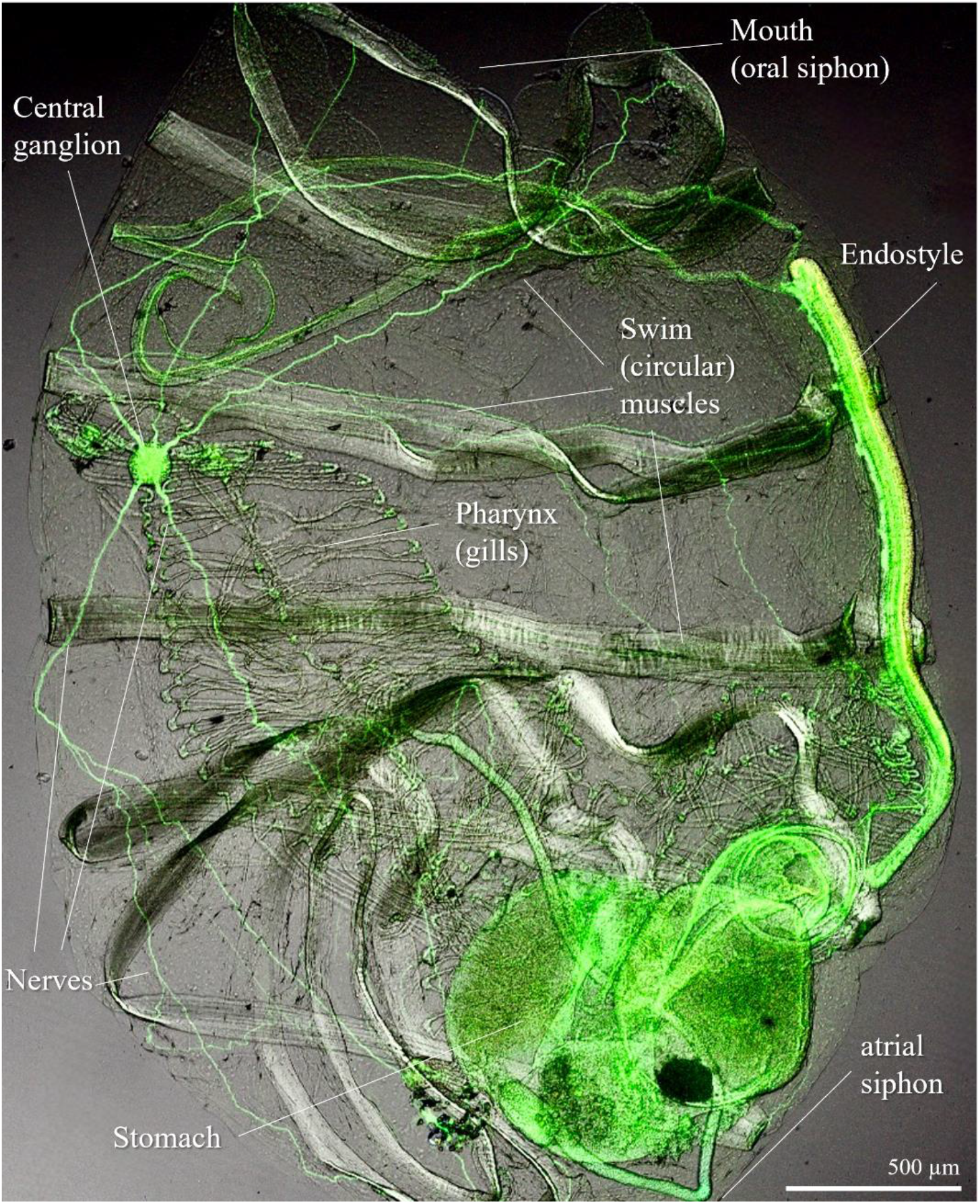
General morphology of *Doliolum* sp. Green labeling indicates anti-tubulin immunoreactivity (see methods for details).

Using glutamate immunohistochemistry in the pelagic tunicate *Doliolum*, we demonstrate the widespread presence of putative glutamatergic axons across all major nerves of the central ganglion, including motor-like terminals on the circular swim muscles in the body wall. In contrast, we did not find GABA-ir neurons in the CNS. These observations suggest that glutamate is a major neurotransmitter in pelagic tunicates and may have neuromuscular functions.

## 2. MATERIALS AND METHODS

Pelagic specimens of *Doliolum* sp. (Chordata, Thaliacea, Doliolida, Fig. 1) were collected in the open ocean of the Northwest Pacific between Haida Gwaii and Vancouver Island, then immediately fixed overnight (10-12 hours) in 4% paraformaldehyde in 0.1 M phosphate-buffered saline (PBS) at +5°C. The fixed animals were washed for 2 hours in PBS and pre-incubated overnight in a blocking solution of 6% goat serum in PBS containing 0.02% Triton X-100 (PBT). The samples were then incubated for 48 hours at +5°C with primary antibodies diluted in 6% goat serum in PBS at a final dilution of 1:100. To label the nervous system with anti-tubulin immunoreactivity, we used the rat monoclonal antibody (AbD Serotec Cat# MCA77G, RRID: AB_325003), which recognizes the alpha subunit of tubulin and specifically binds tyrosylated tubulin (Wehland & Willingham, 1983; Wehland et al., 1983). To label glutamatergic structures, we used a polyclonal anti-Glutamate antibody produced in rabbit (Sigma-Aldrich, Cat# G6642, RRID: AB_259946) at a final dilution of 1:100. For gamma-aminobutyric acid (GABA) immunoreactivity, we used a polyclonal anti-GABA antibody produced in rabbit (Sigma-Aldrich Cat# A2052, RRID: AB_477652) at a final dilution of 1:100.

After a series of PBS washes over 6-8 hours, the animals were incubated for 12 hours with the secondary antibodies. For the anti-tubulin primary antibody, incubation was performed with goat antirat IgG secondary antibodies conjugated to Alexa Fluor 488 (Molecular Probes, Invitrogen, Cat# A11006, RRID: AB_141373) at a final dilution of 1:50. For the anti-Glutamate and anti-GABA primary antibodies, incubation was performed with the goat anti-rabbit IgG cross-adsorbed secondary antibody conjugated to Alexa Fluor 568 (ThermoFisher, Catalog# A-11011, RRID: AB_143157) at a final dilution of 1:50.

To stain the nuclei, tissues were mounted in VECTASHIELD Plus Antifade mounting medium with DAPI (Cat# H-2000). Some of the larger adult tissues were also mounted in Fluorescent Mounting Media (KPL) on glass microscope slides. Slides were viewed on a Nikon Research Microscope Eclipse E800 with Epi-fluorescence using standard TRITC and FITC filters and recorded on a Nikon C1 Laser Scanning confocal microscope.

### Antibody specificity

The rat monoclonal alpha-tubulin antibody used in this study has been extensively characterized, with details on antibody specificity and relevant assays (Wehland & Willingham, 1983; Wehland, Willingham, & Sandoval, 1983). Equally important, this monoclonal antibody has been successfully used in nine ctenophore species (Norekian & Moroz, 2020a), the hydrozoan jellyfish *Aglantha digitale* (Norekian & Moroz, 2020b), and the siphonophore *Nanomia bijuga* (Norekian & Moroz, 2026). To test immunostaining specificity, we omitted either the primary or the secondary antibodies from the procedure. In both cases, we detected no labeling. The anti-Glutamate and anti-GABA antibodies were both produced in rabbit and required the same secondary antibody. Thus, the secondary antibody, all the solutions, and staining protocols remained the same; the only difference was the primary antibodies. The anti-GABA antibody showed no labeling in the nervous system, whereas the anti-Glutamate antibody produced a clear pattern of staining, serving as a good control.

## 3. RESULTS

*Doliolum* is a barrel-shaped, planktonic tunicate, and we used 1-2 cm individuals. The nervous system, visualized with an anti-tubulin antibody (n=8), consisted of a single small ganglion about 150 µm in diameter, located near the dorsal body wall, slightly closer to the oral aperture (**Fig. 2**). Five major nerves exited the ganglion (Fig. 1b, c). Three nerves projected toward the oral aperture, while two others ran toward the atrial opening. There were also four or five very thin nerves. After exiting the ganglion, the major nerves branched extensively, covering the entire body. Some branches targeted the bands of muscles that encircle the body wall (**Fig. 2a, d**). These muscles are responsible for body contraction and for expelling water during jet-propulsion swimming.

**Figure 2.**
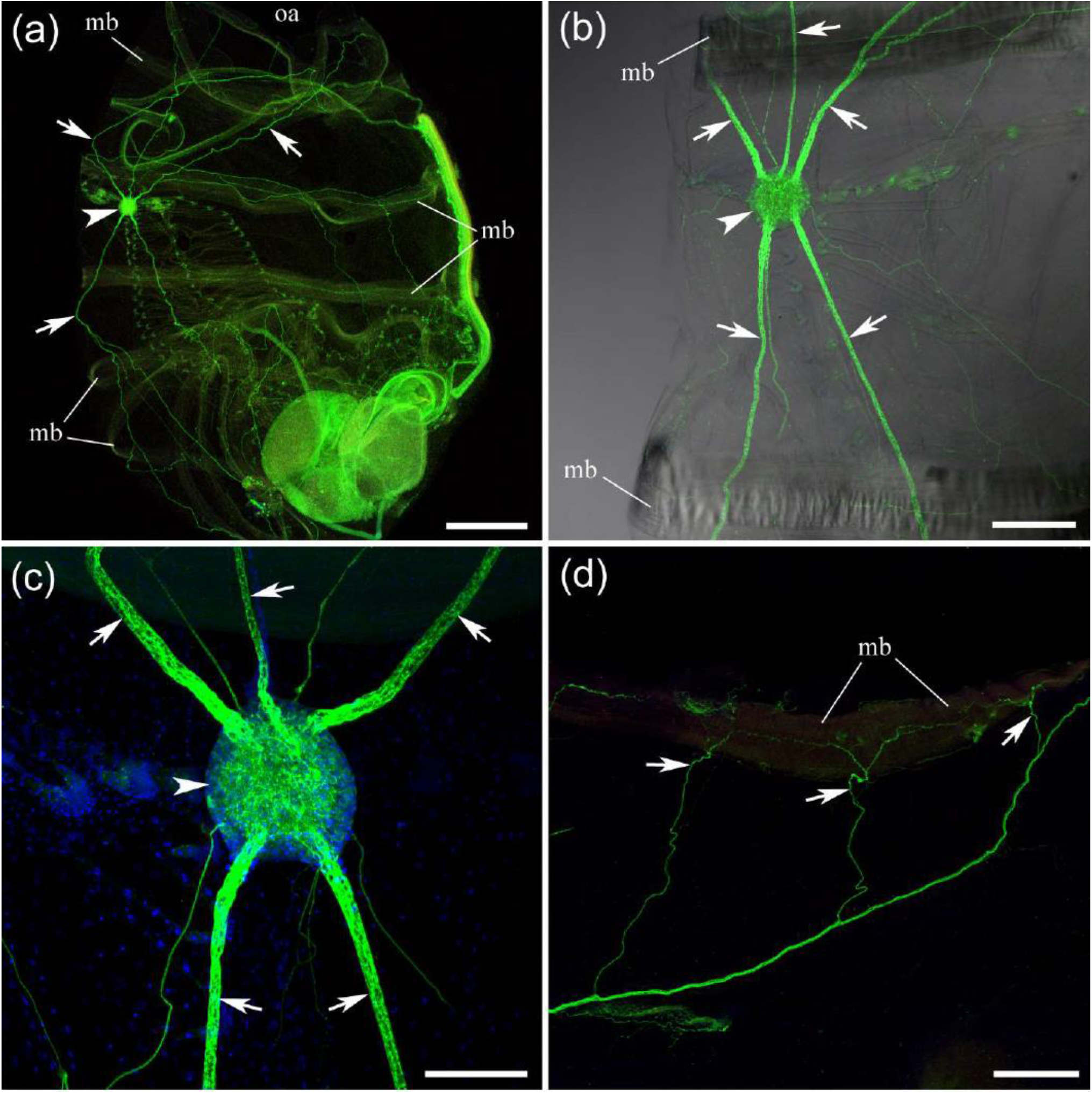
The central nervous system of the pelagic tunicate *Doliolum* labeled with an anti-tubulin antibody (green) and DAPI (blue). (a) - The dorsal ganglion (arrowhead) is located next to the body wall, closer to the oral aperture (oa). Arrows point to some of the major nerves. (b, c) - Five major nerves (arrows) exit the dorsal ganglion (arrowhead): three toward the oral aperture and two toward the atrial opening. Note also about 4-5 thin, smaller nerves produced by the dorsal ganglion. (d) - The nerves branch extensively, with many of the thinner branches (arrows) innervating the circular muscle bands (mb). Abbreviations: oa - oral aperture, mb - muscle band. Scale bars: a - 500 µm, b, d - 200 µm, c - 100 µm.

### Glutamate immunoreactivity

Glutamate immunostaining was confined to the nervous system in all specimens (n=10; **Fig. 3a**). Glutamate-ir signal was also observed in the muscle bands. All five major nerves exiting the central ganglion were glutamate immunoreactive, as were the thin nerves (**Fig. 3c**).

**Figure 3.**
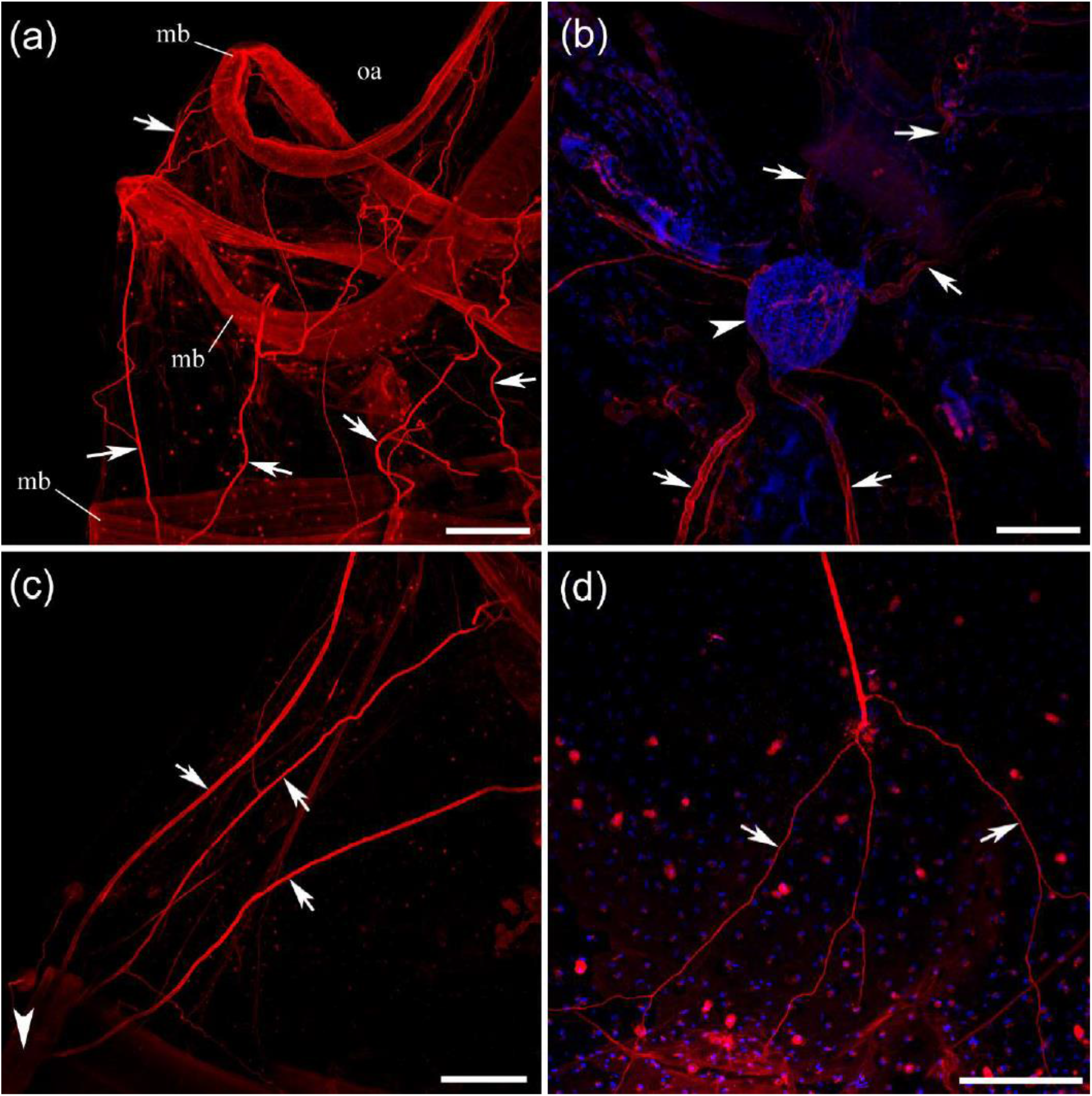
Anti-glutamate immunoreactivity in *Doliolum* (red, with DAPI in blue). (a) - All major nerves and their branches (arrows) are brightly labeled with the anti-glutamate AB. Note that circular muscle bands (mb) also show glutamate-ir signal. (b) - The dorsal ganglion itself (arrowhead) and the initial short stretches of the five major nerves (arrows) do not show significant glutamate-ir staining. (c) - A little further from the dorsal ganglion (arrowhead), the major nerves (arrows) begin to show very bright glutamate-ir labeling. (d) - Such bright glutamate-ir staining is observed throughout the thinnest branches (arrows) and their endings. Abbreviations: *oa* - oral aperture, *mb* - muscle band. Scale bars: a, c - 200 µm, b, d - 100 µm.

However, the dorsal ganglion itself and the very proximal sections of neurites in the neuropile showed very little to no immunoreactivity in all preparations (n=10; **Fig. 3b**). A little farther from the ganglion, glutamate immunoreactivity became very strong (**Fig. 3c**). One possible explanation is poor AB penetration through the ganglion sheaths. However, the anti-tubulin AB didn’t have the same issue. It appears more likely that glutamate was not accumulating in the neuronal cell bodies and was present in detectable quantities only outside the cell bodies - along the neural processes and their terminals. The nerves themselves, their branches, and thin endings were intensely stained (**Fig. 3a, c, d**). All major nerves in this species contain glutamatergic axons.

Circular muscle bands, whose rhythmic contractions pump water through the tunicate body and therefore provide propulsion through seawater, represent one of the main effector organs. We found that many glutamate-ir nerve branches specifically target the circular muscle bundles (**Fig. 4**). Upon reaching the muscle fibers, the nerves branched extensively and made clear contact with them (Fig. 4). The muscle bundles themselves showed low-intensity glutamate IR staining. Notably, glutamatergic neurons in ascidian larvae have sensory-type morphologies (Horie et al., 2008). There is no evidence for the presence of glutamatergic motoneurons in cephalochordates (Candiani et al., 2012), although glutamate concentrations in the neural tube are highest compared to other amino acids (Pascual-Anaya and D’Aniello, 2006), and *in situ* hybridization mapping for vesicular glutamate transporter suggests that at least some glutamatergic cells are likely sensory.

**Figure 4.**
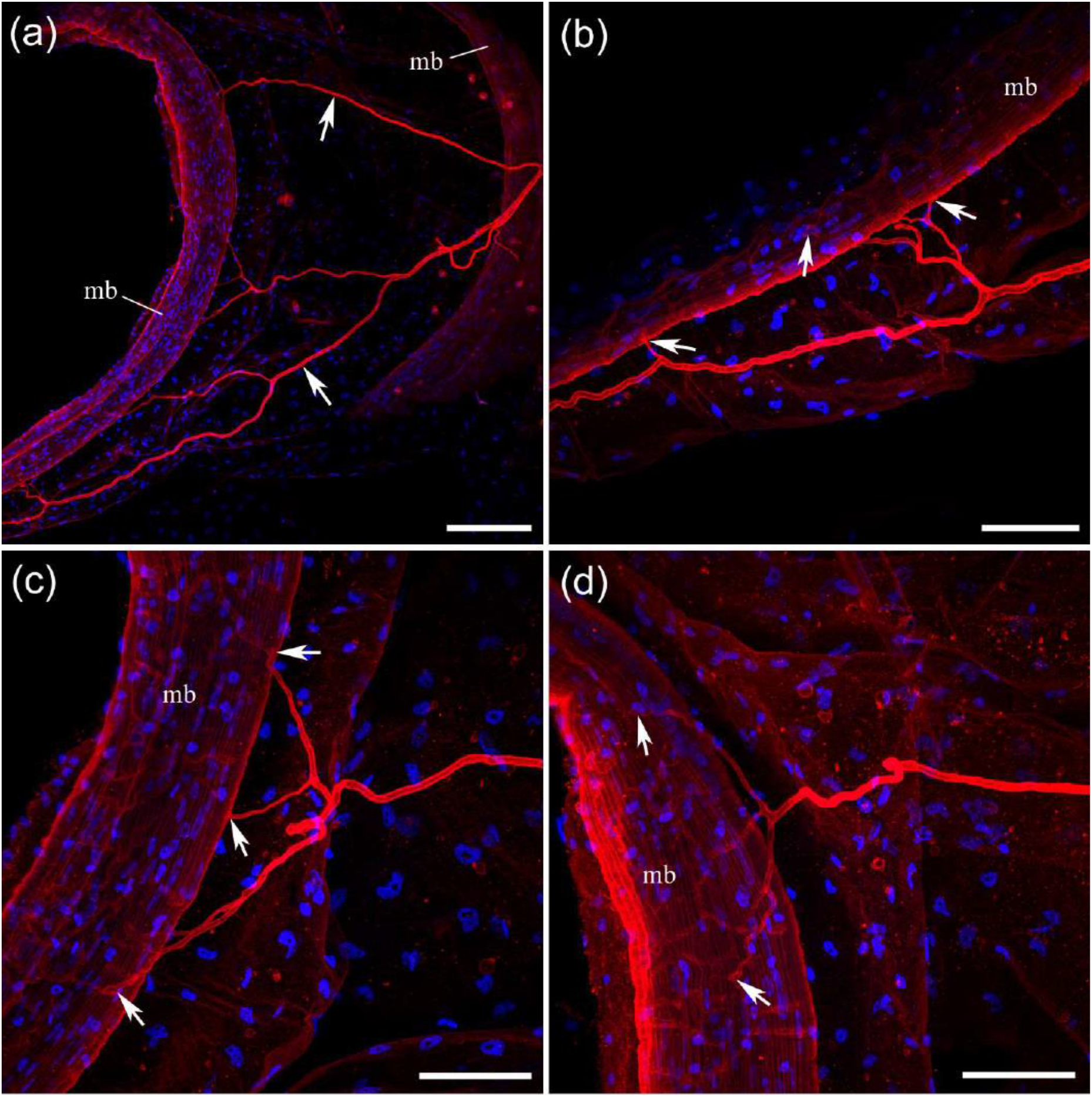
Glutamate-immunoreactive neurites (red) innervate the circular muscle bands in *Doliolum*. DAPI is blue. (a) - Many glutamate-ir nerve branches (arrows) target the muscle bands (mb). (b, c, d) - As glutamate-ir neurites approach the muscle bands, they branch extensively and make clear contact with the muscle (arrows). Note that the muscle bands themselves show a low-intensity glutamate-ir signal. Scale bars: a - 100 µm; b, c, d - 40 µm.

### GABA immunoreactivity

Glutamate and GABA metabolic cycles are coupled, with glutamate serving as a precursor to GABA (Anderson, 2025). Therefore, it was interesting to compare GABA IR and glutamate IR. GABA antibody showed no detectable labeling in the nervous system of *Doliolum*, including the lack of GABA-ir signal in central ganglion, major nerves, smaller branches, and their endings (n=4; **Fig. 5**). The same lot of GABA AB was successfully used during the same time period on two molluscan species (*Clione* and *Melibe*, not shown) as a control, proving that the AB were working.

**Figure 5.**
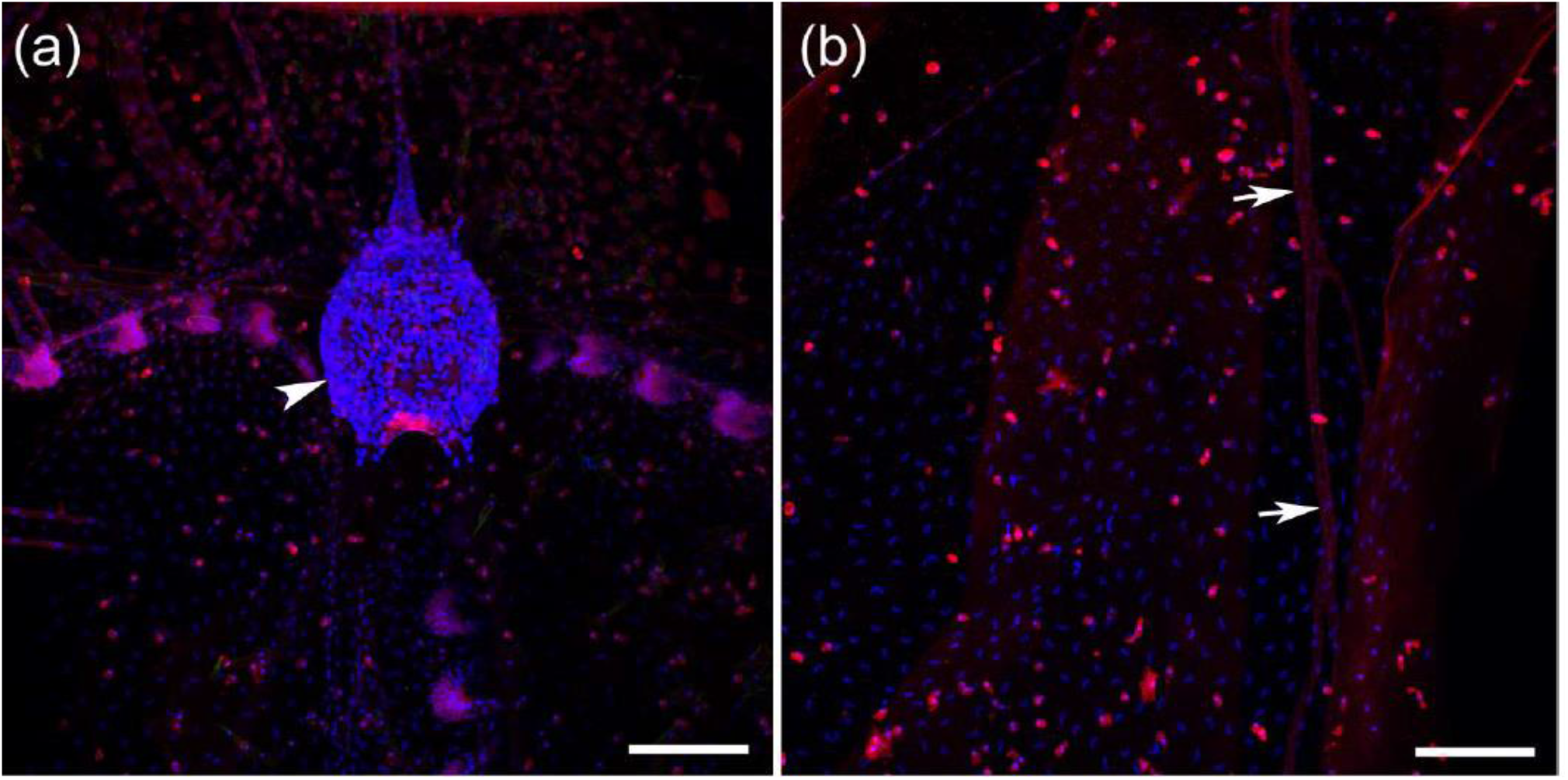
Anti-GABA immunoreactivity in *Doliolum* (red, DAPI is blue). (a) - The dorsal ganglion (arrowhead) shows no GABA ir. Several hundred small DAPI-stained nuclei are present within the ganglion. (b) - Arrows indicate the location of a major nerve at a significant distance from the dorsal ganglion. GABA- ir cells of unknown identity were also noted. The nerve shows no GABA-ir labeling. Scale bars: 100 µm.

## 4. DISCUSSION

In the animal phyla studied, we may observe alternative selection of glutamate-mediated transmitter chemistry across different components of neural circuits (Moroz et al., 2021). In insects, glutamate is recruited predominantly by motor neurons to control skeletal muscles, but by sensory cells and interneurons in vertebrates. In vertebrates and ascidians (Manni and Pennati, 2016), acetylcholine is the classical neuromuscular transmitter, whereas in insects, acetylcholine is a primary sensory neurotransmitter.

The most surprising finding of this study was the high abundance of glutamatergic neural processes in *Doliolum*, particularly in the peripheral nerves, where axonal-like terminals innervate the body wall muscles. This innervation pattern (with putative neuromuscular junctions similar to those described by Bone, 1959) suggests that glutamate serves as a neuromuscular transmitter. In other words, the glutamatergic system in *Doliolum* can be motor, as in insects, but not as in ascidians (Horie et al., 2008; Takamura et al., 2010), cephalochordates (Candiani et al., 2012), and vertebrates. The hypothesis of a neuromuscular role for glutamate in pelagic tunicates should be tested experimentally, ideally by detecting glutamate release, analyzing its role in muscle contractility, and identifying specific glutamate receptors. Nevertheless, we should not completely exclude glutamate as a sensory transmitter (e.g., in proprioceptive pathways). Further studies of glutamate’s possible role as a neuromuscular transmitter would be of great evolutionary significance, especially given the early divergence of the chordate lineage within the bilaterian phylogeny (Serra Silva et al., 2025).

The role of glutamate as the evolutionary earliest neuromuscular transmitter was proposed based on electrophysiological and pharmacological data on ctenophores, descendants of the earliest-branching animal lineage with independently evolved neural systems (Moroz et al., 2014; Moroz et al., 2021). Glutamatergic neurons were recently identified in the ctenophore subepithelial nets (Moroz and Norekian, 2026a) and in pelagic hydrozoans (Moroz and Norekian, 2026b) using the same antibodies. Equally important is our observation of Glutamate IR in muscles of representatives across three metazoan lineages: ctenophores (Moroz and Norekian, 2026a), cnidarians (Moroz and Norekian, 2026b), and in this study. Such parallel recruitment of glutamate in specific muscle and neural populations might be associated with an anaplerotic role for both glutamate and GABA, further highlighting the deep metabolic roots of neuronal origins from secretory cells and respective neurotransmitters repeatedly recruited as energy-relevant signaling molecules for costly stress/injury adaptive responses (Moroz, 2009, 2014, 2021).

The second surprise in the current study was that we were unable to detect GABAergic neurons in *Doliolum*, although these types of neurons are abundant in both amphioxus and the ascidian *Ciona*, (Zega et al., 2008). Thus, glutamatergic innervation appears to be widely distributed in *Doliolum sp*., while GABAergic elements are not.

In sum, these morphological observations call for systematic functional and pharmacological studies of pelagic tunicates beyond glutamate and GABA, taking advantage of their astonishing diversification of forms and functions (Bone, 1998), ecologies, and evolutionary significance.

## Acknowledgments

We thank FHL for its excellent facilities, including the Nikon Laser-Scanning confocal microscope. This research was supported by the National Science Foundation grant (IOS-2341882) and National Institutes of Health grant (5R01NS11449) to LLM.

## Conflict of interest

None of the authors has any known or potential conflict of interest, including any financial, personal, or other relationships with other people or organizations within three years of beginning the study that could inappropriately influence, or be perceived to influence, their work.

## Role of the authors

All authors have full access to all study data and take responsibility for data integrity and the accuracy of the data analysis. Research design: LLM, TPN. Acquisition of data: TPN. Analysis and interpretation of data: LLM, TPN. Drafting of the article: TPN, LLM. Funding: LLM.

## Notes

### Competing Interest Statement

The authors have declared no competing interest.

